# Micromechanical homogenisation of a hydrogel-filled electrospun scaffold for tissue-engineered epicardial patching of the infarcted heart

**DOI:** 10.1101/2020.11.08.373209

**Authors:** Tamer Abdalrahman, Nicolas Mandel, Kevin L. Sack, Nicola M. Pugno, Deon Bezuidenhout, Georges Limbert, Francesco Moscato, Neil H. Davies, Thomas Franz

**Affiliations:** Division of Biomedical Engineering, University of Cape Town, Cape Town, South Africa; Julius Wolff Institut, Charité – Universitätsmedizin Berlin, Berlin, Germany; University of Applied Sciences, Vienna, Austria; Department of Surgery, University of California at San Francisco, San Francisco, California, USA; Laboratory of Bio-inspired & Graphene Nanomechanics, Department of Civil, Environmental and Mechanical Engineering, University of Trento, Trento, Italy; School of Engineering and Materials Science, Queen Mary University, London, UK; Ket-Lab, Edoardo Amaldi Foundation, Italian Space Agency, Rome, Italy; Cardiovascular Research Unit, University of Cape Town, Cape Town, South Africa; Bioengineering Science Research Group, University of Southampton, Southampton, UK; Center for Medical Physics and Biomedical Engineering, Medical University of Vienna, Austria

**Keywords:** Fibrous scaffold, myocardial infarction, mean field homogenisation, foam mechanics, finite element method, composite material

## Abstract

This study aimed at developing a formulation to link microscopic structure and macroscopic mechanics of a fibrous scaffold filled with a hydrogel for use as a tissue-engineered patch for local epicardial support of the infarcted heart. Mori-Tanaka mean field homogenisation, closed-cell foam mechanics and finite element (FE) methods were used to represent the macroscopic elastic modulus of the filled fibrous scaffold. The homogenised constitutive description of the scaffold was implemented for an epicardial patch in a FE model of a human cardiac left ventricle (LV) to assess effects of patching on myocardial mechanics and ventricular function in presences of an infarct. The macroscopic elastic modulus of the scaffold was predicted to be 0.287 MPa with the FE method and 0.290 MPa with the closed-cell model for the realistic fibre structure of the scaffold, and 0.108 and 0.540 MPa with mean field homogenization for randomly oriented and completely aligned fibres. Epicardial patching was predicted to reduce maximum myocardial stress in the infarcted LV from 19 kPa (no patch) to 9.5 kPa (patch), and to increase the ventricular ejection fraction from 40% (no patch) to 43% (patch). The predictions of the macroscopic elastic modulus of the realistic scaffold with the FE and the closed-cell model agreed well, and were bound by the mean field homogenisation prediction for random and fully aligned fibre orientation of the scaffold. This study demonstrates the feasibility of homogenization techniques to represent complex multiscale structural features in an simplified but meaningful manner.

## 1. Introduction

Cardiovascular disease (CVD) remains the leading cause of death worldwide. In 2013, roughly 30% of all deaths in the United States of America could be attributed to cardiovascular disease [1]. Myocardial infarction (MI), one of the biggest contributors to CVD, is caused by a blockage of the blood supply to the myocardium that initiates ischemia and subsequent tissue death. Post-infarct treatment concepts aim at halting the degenerative progress. The increasing distention of the left ventricle induces induce cellular processes that in lead to stronger degeneration. Intramyocardial biomaterial injection and epicardial patching are two promising approaches to counteract the deleterious effects of ventricular distention by altering the underlying mechanics of the heart [2], [3].

Both patching and injection therapy can benefit impaired myocardium in two ways. The first is the delivery of bioactive compounds into the myocardium. These compounds influence the patho-biological pathways mobilized by MI and intervene either through paracrine signalling or through delivery of active growth factors [2]–[5]. The second is the mechanical support that biomaterial injectates and epicardial patches can provide to the impaired myocardium [2], [5], [6], which has been shown to decrease adverse remodelling [7].

Patching treatment concepts have recently grown in significance due to improved cell retention capabilities, relative ease in which they deliver bioactive molecules, substantial mechanical support [5] and as patches can be easily tailored to a specific size and shape. Patches are almost exclusively made up of scaffold structures consisting of interconnecting fibrous networks or porous structure. Scaffold characteristics depend on the method of creation (e.g. electrospinning, 3D printing [8]), the underlying material choice (e.g. polymer, decellularised matrices [9]) and the length scales of the structure (i.e. fibre diameter and pore size). This results in diverse structural properties of scaffolds on both the macro- and micro-scale.

The combination of electrospun fibrous scaffolds and hydrogels has been studied previously [10]–[15] and is a promising treatment concept, exploiting the advantages that each material offers. Different structures and mechanical properties from both groups of materials in combination with different fabrication techniques can lead to a large variety in compound materials [16]. The mechanical characterization of these combined materials is scarcely described. Kundanati et al. [17] obtained the elastic modulus of a silk scaffold filled with silk hydrogel matrix in compression tests and compared the results to analytical data from combination of open-cell and closed cell models [18]. Their results showed an increase in elastic modulus of the composite compared to either of the basic structural components, and experimental and analytical results were similar. Strange et al. [19] determined the elastic modulus of a polycaprolactone electrospun scaffold filled with alginate hydrogels of different concentrations in indentation and tensile tests. They observed strain toughening behaviour similar to that occurring in conventional composite materials, but also found large differences between the two test methods.

This study aimed at linking microscopic and macroscopic mechanical description of a hydrogel-filled electrospun polymeric scaffold and exploring the use of such scaffolds as epicardial patches for treatment of MI. We introduce a combination of analytical and numerical models to derive a homogenized macroscopic description of the scaffold mechanics of these composite materials and, identify the effects of the scaffold as epicardial patch on left ventricular mechanics and function in the infarcted heart.

## 2. Materials and Methods

This study is divided into two parts: (I) Determination of the homogenized elastic modulus of an electrospun hydrogel-filled scaffold using a finite element (FE) model and comparison to analytical models. (II) Investigation of the treatment of an acute left ventricular (LV) infarct with an epicardial electrospun hydrogel-filled scaffold patch utilizing the FE homogenization method.

### 2.1 Electrospun scaffold

Pellethane© 2363–80AE (Dow Chemicals, Midland, MI, USA), a medical grade aromatic poly(ether urethane) with a hard segment consisting of 4,4-methylenebis(phenylisocyanate) and 1,4-butanediol and a soft segment of poly(tetramethylene oxide) glycol (Mn = 1000 g/mole) [20] was used.

A 15 wt% Pellethane© solution was obtained by dissolving 8 g of Pellethane© pellets in 53.3 g of tetrahydrofuran (THF, source) at 37°C for 8 hours. Using a custom-made rig, the Pellethane® solution was electro-spun from a hypodermic needle (SE400B syringe pump, Fresenius, Bad Homburg, Germany) onto a rotating and bi-directionally translating tubular target (hypodermic tubing, Small Parts, Loganport, IN, USA). The spinning process parameters were, for subcutaneous and circulatory samples, respectively: solution flow rate: 6 and 4.8 mL/h, target outer diameter: 25.0 and 2.2 mm, target rotating speed 4400 and 9600 rpm, target translational speed: 2.6 mm/min; potential difference: 13 and 15 kV; source-target distance: 250 mm.

### 2.2 Geometrical Modelling

#### 2.2.1 Electrospun scaffold

Micro-computed tomography (μCT) images of the electrospun fibrous scaffold obtained at the University of Southampton were processed with the Simpleware ScanIP (Synopsys, Mountain View, CA, USA), see Figure 1(a). Spatial parameters of the μCT images are provided in Table 1. A region of the scaffold sample was selected to provide a representative volume element (RVE) of the interconnected fibre structure of the scaffold. Fibre surfaces were smoothed, and any residual islands of fibres were excluded. The mean fibre diameter was 6.7 μm. The reconstructed geometry of the scaffold RVE is presented in Figure 1(b). To simulate complete filling of the fibrous scaffold with hydrogel, void space between the fibres of the scaffold RVE was identified in Simpleware by inverting the scaffold mask and treated as hydrogel domain, as illustrated in Figure 1(c).

**Figure 1.**
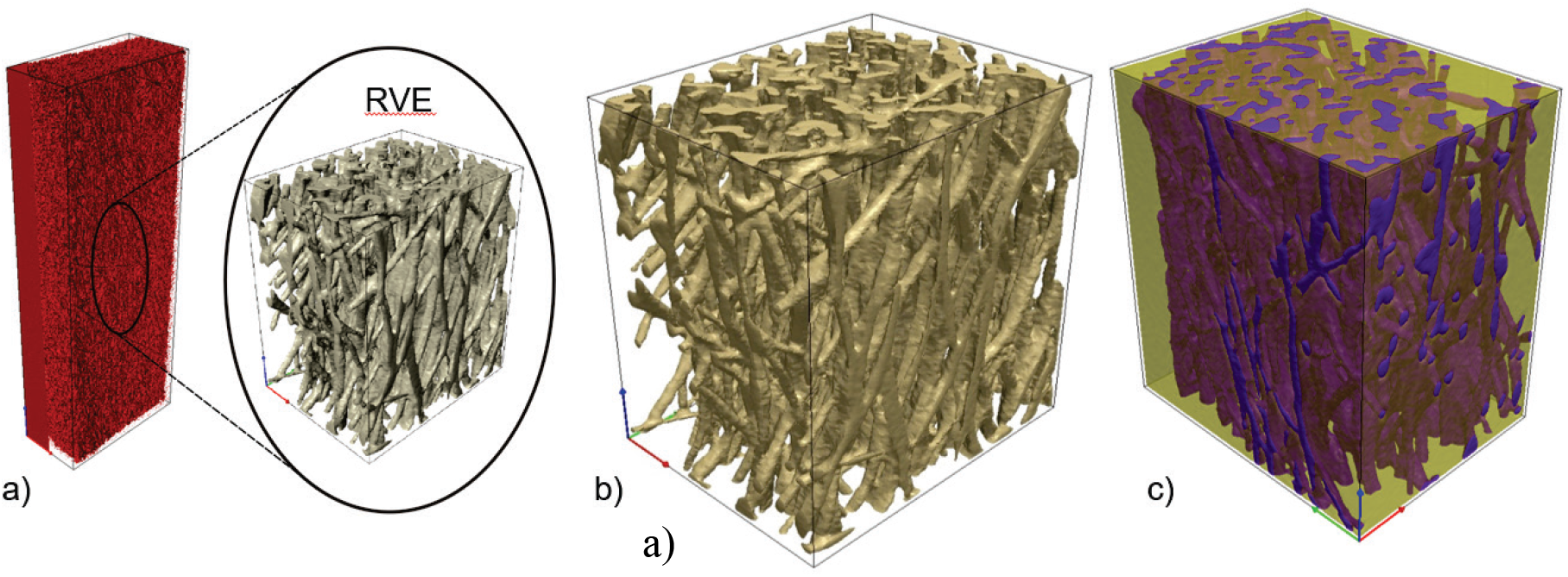
a: MicroCT representation of electrospun scaffold sample and smaller representative volume element (RVE). b: 3D reconstructed geometry of representative volume element of electrospun scaffold. c: Electrospun scaffold (purple) filled with hydrogel (beige, opacity = 70%).

**Table 1.** Basic spatial properties of scaffold μCT data

| Axis | Image Extent (mm) | Spacing (mm) | Voxel Extent |
| --- | --- | --- | --- |
| X | 0.2226 | 0.0021 | 106 |
| Y | 0.2877 | 0.0021 | 137 |
| Z | 0.3213 | 0.0021 | 153 |

#### 2.2.2 Cardiac left ventricle and epicardial patch

The geometry of a left ventricle was reconstructed from a normal human subject, utilising the data from Genet et al. [21] An acute infarct in the LV free wall was simulated in the anterior wall, midway between the base and the apex of the heart. Infarction was assumed to affect tissue within a radius of 25 mm, resulting in a fully transmural infarct. For the treated infarct case with an epicardial patch, a layer with a thickness of 2.0 mm was added to the epicardial ventricular surface in the infarct region (Figure 2).

**Figure 2.**
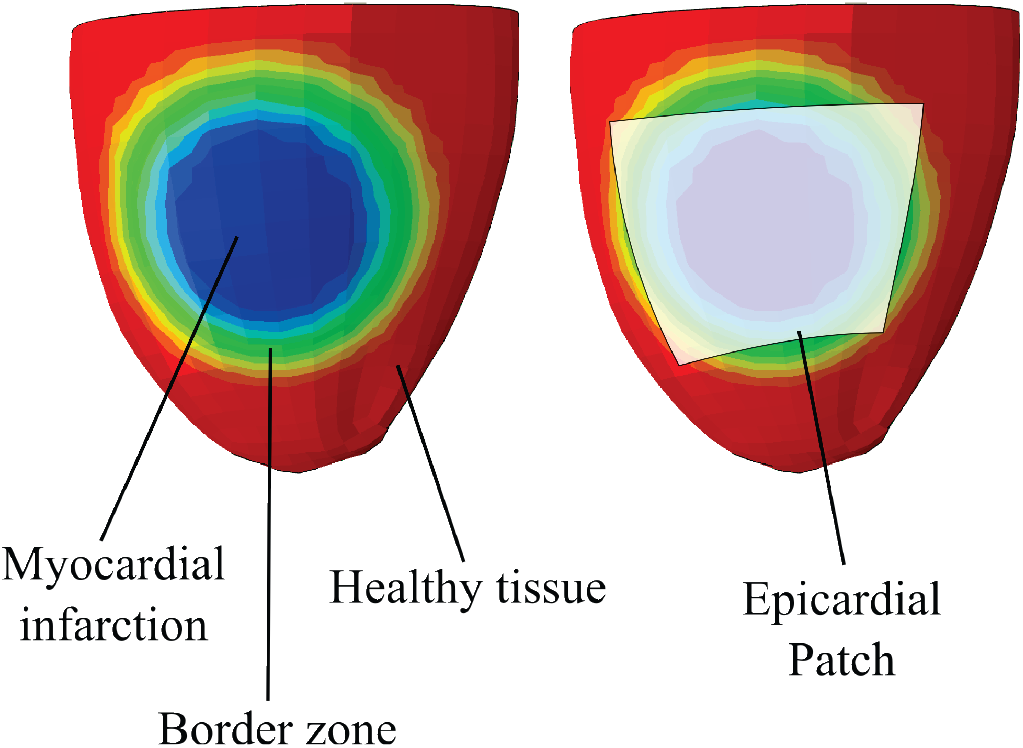
The application of an idealized patch to treat myocardial infarction improves function and regional stress within the heart. Schematic illustration of an LV showing the description of myocardial infarction (MI) and the placement of the patch.

### 2.3 Finite Element Modelling

#### 2.3.1 Mesh generation

##### a) Scaffold

The scaffold, and hydrogel were meshed by using Simpleware ScanIP (Synopsys, Mountain View, CA, USA). Continuous transitions were used between scaffold fibres and hydrogel, with mesh refinement near fibre-hydrogel interfaces, see Figure 3. The RVE comprised scaffold fibres with a volume fraction of 21.6% and 364,429 elements and hydrogel with a volume fraction of 78.4% and 524,580 elements. The meshed geometry was imported into Abaqus Standard (Dassault Systèmes, Johnston, RI, USA).

**Figure 3.**
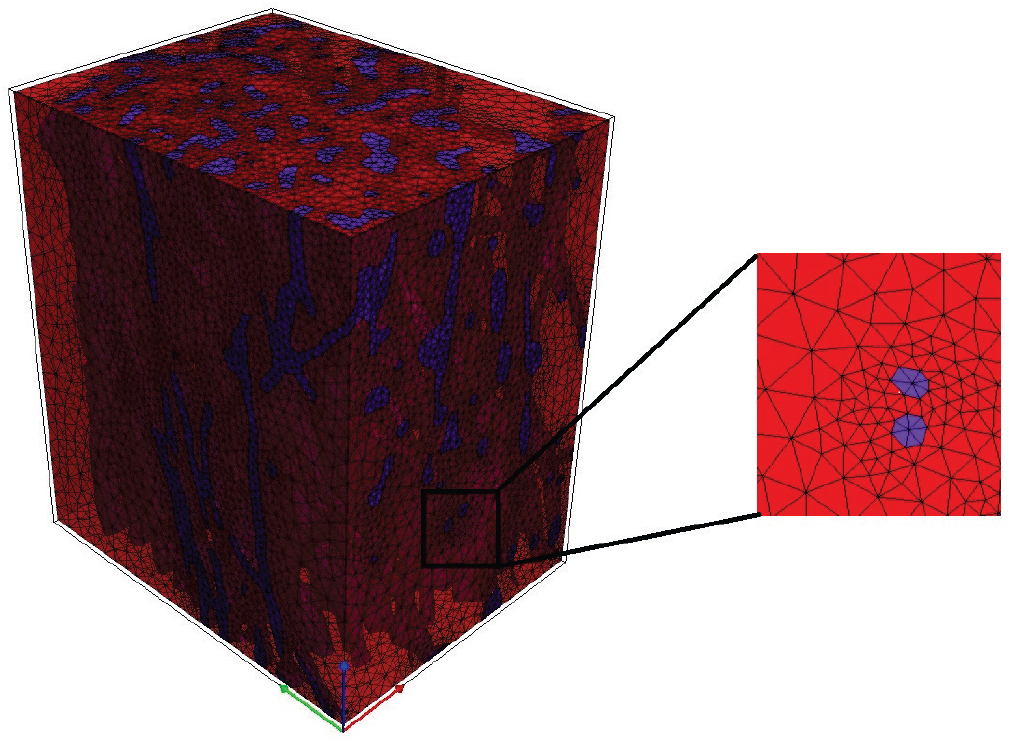
Scaffold and hydrogel mesh as exported by ScanIP with a magnification as an example of mesh refinement results

##### b) Cardiac left ventricle and homogenized patch

LV geometry was meshed using approximately 3,500 linear hexahedral elements (C3D8H). The homogenized patch was meshed using 72 quadrilateral shell elements (S4R, arranged in a 9 x 8 grid, using the underlying hexahedral element faces).

#### 2.3.2 Constitutive laws

##### a) Scaffold and hydrogel

The scaffold fibres and hydrogel were represented with a Neo-Hookean strain energy density function *ψ* [20], [22], [23] defined as:

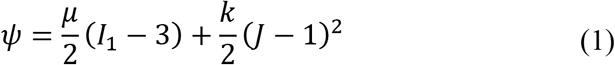

where *μ* and *k* are the ground state shear and bulk modulus respectively. *I*_1_ and *J* are the first and third invariant of right Cauchy-Green deformation tensor, **B.** The latter is given by:

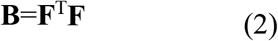

where **F** is the deformation gradient tensor [24]. All material parameters are provided in Table 2.

**Table 2.** Mechanical properties of scaffold material and hydrogel

| Components | Constitutive law | Parameters |
| --- | --- | --- |
| Scaffold fibres | Hyperelastic | $\mu=0.96$ MPa, $k=2.08$ MPa [25] |
| Hydrogel | Hyperelastic | $\mu=0.35$ KPa, $k=3.33$ KPa [26] |

##### b) Healthy and infarcted cardiac left ventricle

The passive mechanics of cardiac tissue were represented following Holzapfel and Ogden [27]. The isochoric and volumetric responses are governed by the strain energy potentials

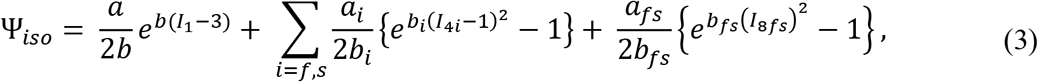

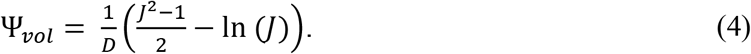

Equation (3) is defined through eight material parameters *a, b, a*_*f*_, *b*_*f*_, *a*_*s*_, *b*_*s*_, *a*_*fs*_, *b*_*fs*_ and four strain invariants that deal with isotropic deformation, *I_1_*, deformation in the fibre and sheet direction, *I_4f_* and *I_4s_* respectively, and shear deformation that couples the fibre-sheet plane, *I_8fs_*.

Equation (4) is defined through *J* and a penalty term *D,* which is inversely proportional to the bulk modulus (*D* = 2/*K*) and therefore characterises the degree of compressibility. For a deformation that perfectly volume, *J* =1.

This passive material model (Eqs. 3 and 4) ensures that the material exhibits the well-documented exponential and anisotropic response to strain [28]–[30] while also enforcing incompressibility.

The description of the time-varying elastance model of active force [31] development is specified as:

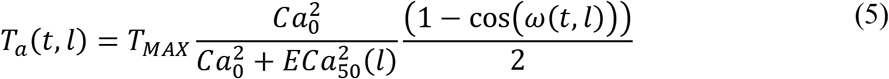

where *T*_*max*_, the maximum allowable active tension, is multiplied with a term governing the calcium concentration, and a term governing the timing of contraction. Both terms depend on sarcomere length *l*, which in turn depends on the strain in the fiber direction. The active tension generated from this representation captures length-dependent effects such as Frank Starling’s Law [32].

The infarcted left ventricle was represented by altering material parameters of the myocardium in the infarct region such that the myocardium was non-contractile and half as stiff as the healthy tissue [33]. A functional border zone (see Figure 2) was implemented through linear transition of the elastic modulus and contractility from the infarct region to the healthy tissue region.

##### c) Homogenized patch

The elastic modulus resulting from homogenization techniques and the assumption of near incompressibility (Poisson’s ratio v = 0.45) and linear elasticity were used to describe the mechanical properties of the patch.

#### 2.3.3 Boundary conditions and loading

##### a) Scaffold model

For study part I, the boundary conditions used are shown in Figure 4. One surface of the RVE was fixed in all directions and the displacement was applied normally on the opposite surface. The remaining sides were free in displacement direction. This generated a uniform field of tensile strain up to 0.1 (λ=1.1).

**Figure 4.**
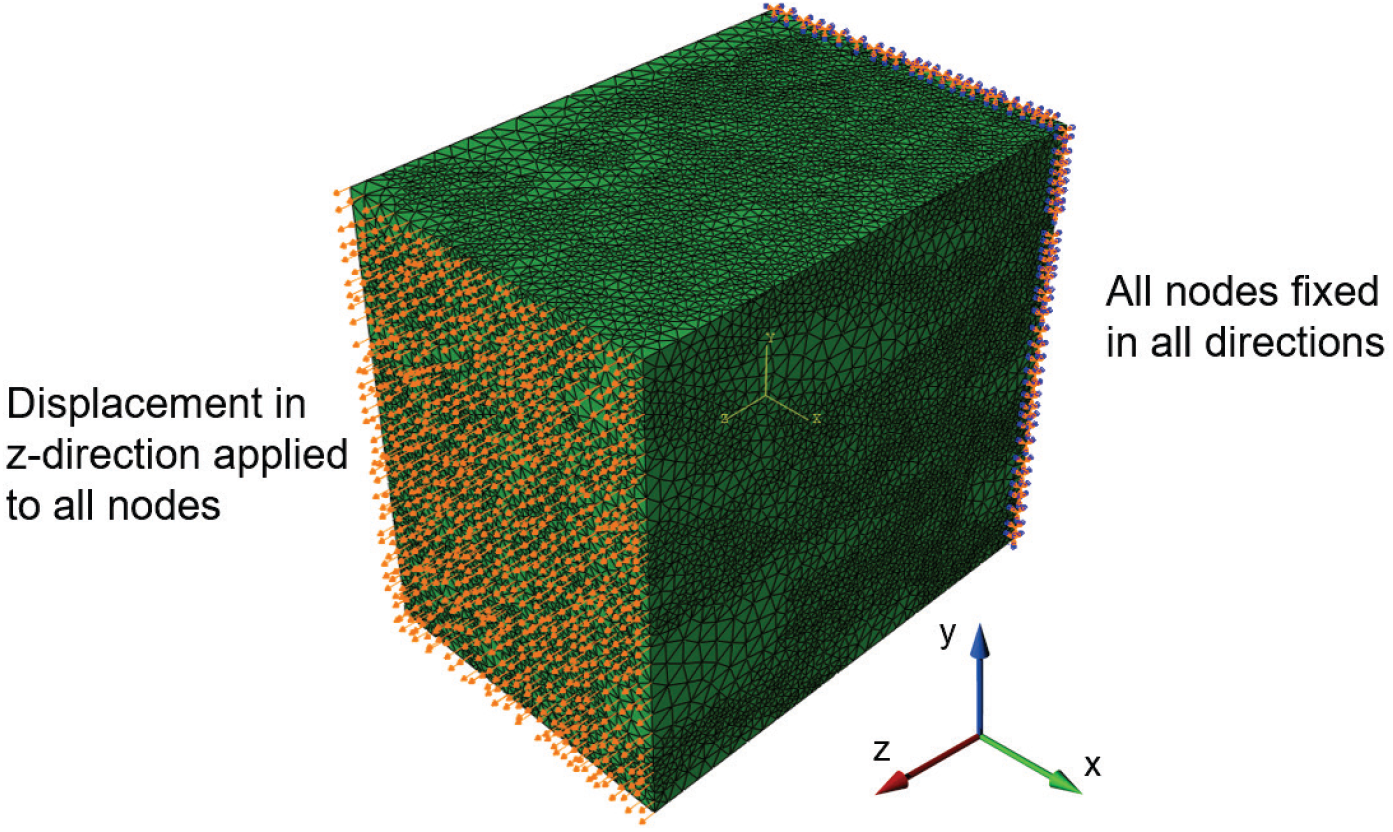
Illustration of boundary conditions and loading of scaffold RVE model.

##### b) Left ventricular model

In the LV model, all nodes on the truncated base were fixed in-plane. The endocardial ring of the LV truncated base was coupled to a fictitious fixed point at its centre using a distributed kinematic constraint to prevent rigid body translation.

The FE model of the LV was coupled to a lumped circulatory model to simulate loading consistent with the physiological beating heart [34]. The circulatory model parameters are fixed across simulations, which ensure the difference cases experience the same pre- and after-load during the cardiac cycle. During diastole the LV endocardial surface experiences gradual loading until 12mmHg is reached. During systole the cavity pressure depends on the force generation of the contracting LV interacting with the lumped circulatory model.

#### 2.3.4 Simulations and analyses

##### a) Scaffold homogenization

Uniaxial stretch corresponding to maximum of 10% global strain was simulated with the FE model of the hydrogel-filled scaffold and resulting von Mises stress, *σ*, and logarithmic maximum principal strain, *ε*, were captured. Nominal strain and normal stress in loading direction were volume-normalized to account for variable element sizes and meshing anomalies (<σ>, <ε>), and used to calculate the homogenised elastic modulus, E, according to

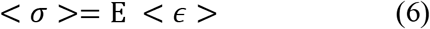

##### b) Left ventricle

Cardiac simulations used the same LV geometry for healthy case, untreated acute MI case, and acute MI case with epicardial patch. Pressure-volume curves and myocardial stresses were captured and used to assess the treatment effects of the epicardial patch.

### 2.4 Analytical model of homogenization

#### 2.4.1 Mean field homogenization

Mean field (MF) homogenization of the mechanical properties of the scaffold-hydrogel composite was undertaken by analysis of the RVE described above. The RVE served as a statistical representation of material properties as shown in Figure 1. It should contain enough information to describe the behaviour of a microstructure [35].

Homogenization techniques, as stated before, have been based often on direct finite element analysis of an RVE at micro scale using macroscopic values as the boundary conditions [36], [37].

In the case of a two-phase composite, the strain field over an RVE comprising inclusion phase and matrix phase are related as:

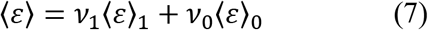

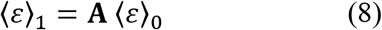

where *v*_i_ is the volume fraction, and subscripts 0 and 1 are identification indices corresponding to the inclusion phase and matric phase respectively. The average strains per each phase are correlated by a strain concentration tensor **A.** Using Eshelby’s solution [38], it can be found that strain inside inclusion (ε) is uniform and related to remote strain (Ω):

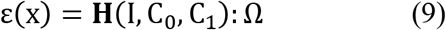

Where C_0_, and C_1_ are the stiffnesses of the inclusion and matrix respectively and **H** is the single inclusion strain concentration tensor define as follows:

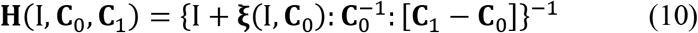

where **ξ**(I,C_0_) is the Eshelby tensor.

The Mori-Tanaka homogenization model, which is used in further computations, assumes that the strain concentration tensor **A** is equal to the strain concentration tensor of the single inclusion problem **H** [38], [39]. Benveniste [40] gives an interpretation of Mori-Tanaka’s model: Each inclusion behaves like an isolated inclusion in the matrix being exhibited to the average matrix strain 〈*ε*〉_0_ as a far-field strain. The effective stiffness can be given by

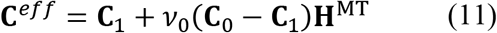

where **H**^MT^ is the Mori-Tanaka strain concentration tensor [37].

For our MF analytical model, we considered two cases: One for completely aligned fibres and the other for randomly oriented fibres.

#### 2.4.2 Foam mechanics

The mechanical behaviour of the fibrous scaffold can be predicted by using the open cell model where the elastic modulus of the fibrous scaffold is obtained by [18]:

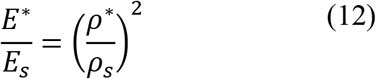

where *E*_s_, *ρ*_s_, *E*\*, and *ρ*\* are elastic modulus and density of the individual fibres and the fibrous scaffold, respectively.

The homogenized elastic modulus of the scaffold filled with hydrogel can be estimated by assuming the cell filled with hydrogel as shown in Figure 5. The homogenized elastic modulus E_hom_ can be calculated using [18]:

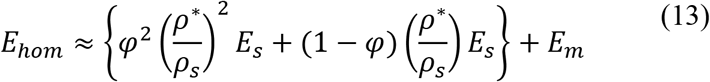

where *φ* is the ratio of volume of edge to total volume of face and E_m_ is the elastic modulus of the hydrogel matrix. The contribution of edge bending and face stretching are represented by the first and the second term, respectively, and the contribution of the hydrogel matrix is represented by the third term.

**Figure 5.**
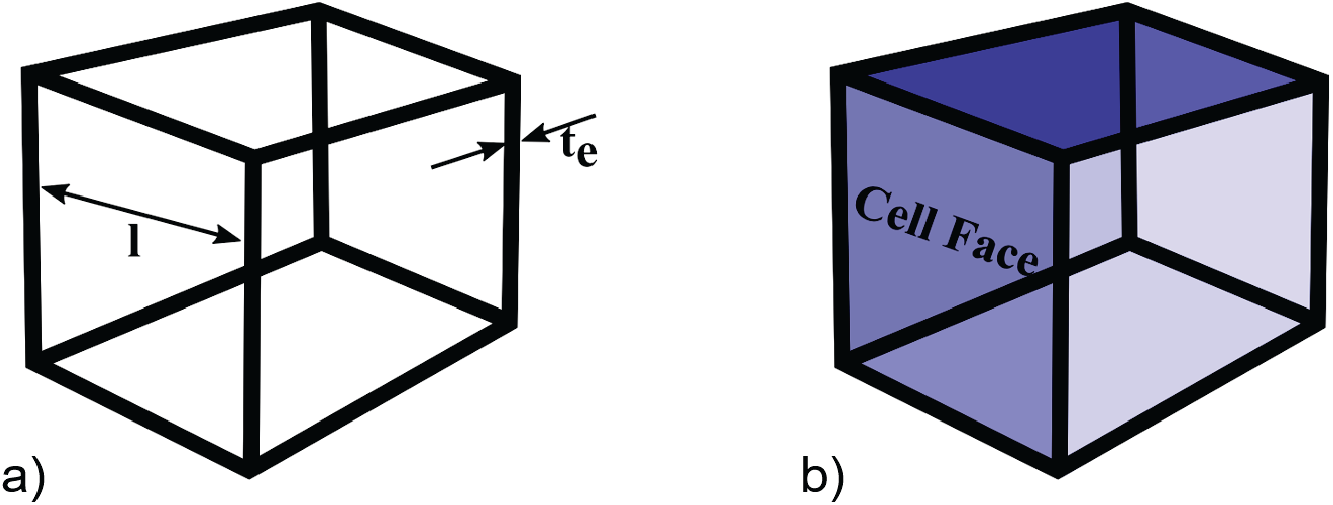
(a) Representative open cell model used for predicting the elastic modulus of scaffold; (b) Corresponding closed cell showing the cell edges and cell face used for predicting the modulus of the composite scaffold.

The homogenization process is based on deformations of the approximated unit cell. In the model of Kundanati et. al [17], the solid fraction of the edges can be estimated with

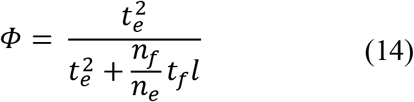

This relationship describes the amount of solid fraction that is contained in compression-bearing edges as the main determinant for mechanical stability, including *n*_*f*_ and *n*_*e*_, which are the number of faces meeting at an edge and the numbers of edges of a face, respectively, and *l* which is the size of single pore [41]. For approximation, we consider *t*_*f*_ and *t*_*e*_ to be equal and of the value of the fibre diameter of 6.7 μm.

## 3. Results

### 3.1 Scaffold homogenization

The strain and stress distributions within the scaffold predicted with the FE model (Figure 6a and b) exhibit significant heterogeneity, with concentrations of stress and strain up to four orders of magnitude higher than the average values. The hydrogel displays strains one order of magnitude higher than the scaffold fibres (Figure 6c), whereas the stress in the hydrogel (Figure 6d) is one order of magnitude smaller than the stress in the scaffold fibres.

**Figure 6.**
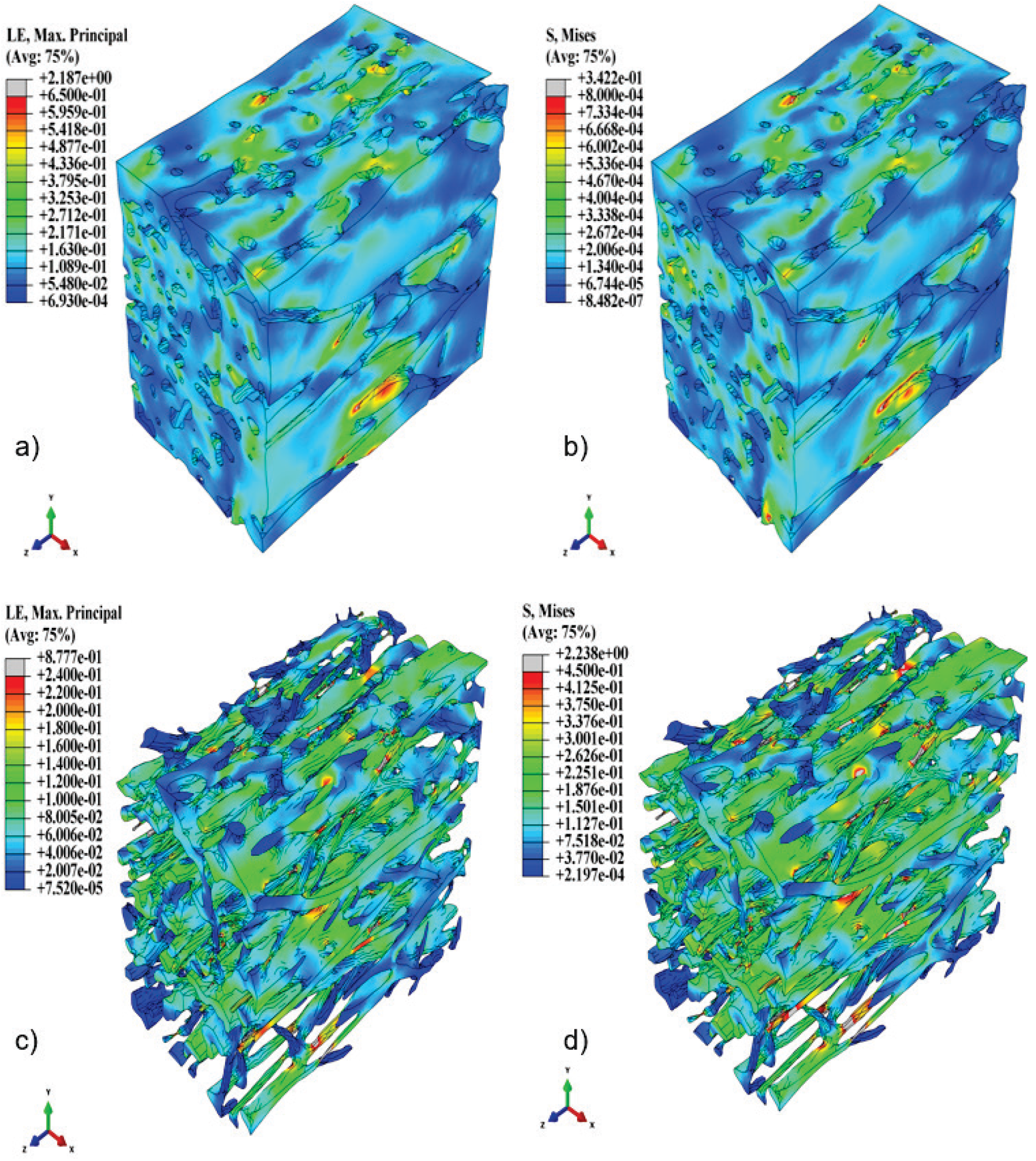
Contour plots of the logarithmic maximum principal strain (a) and von Mises stress (MPa) (b) in the hydrogel, and of the logarithmic maximum principal strain (c) and von Mises stress (MPa) (d) in the scaffold structure.

Figure 7 illustrates the homogenized elastic modulus of the RVE obtained from the FE simulation (E_hom_ = 0.287 MPa), the closed cell model derivation (E_hom_ = 0.29 MPa), and the mean field homogenization for completely aligned and randomly oriented fibres (E_hom_ = 0.540 MPa and 0.108 MPa, respectively). The elastic modulus obtained with the closed cell model agrees well with the FE results.

**Figure 7.**
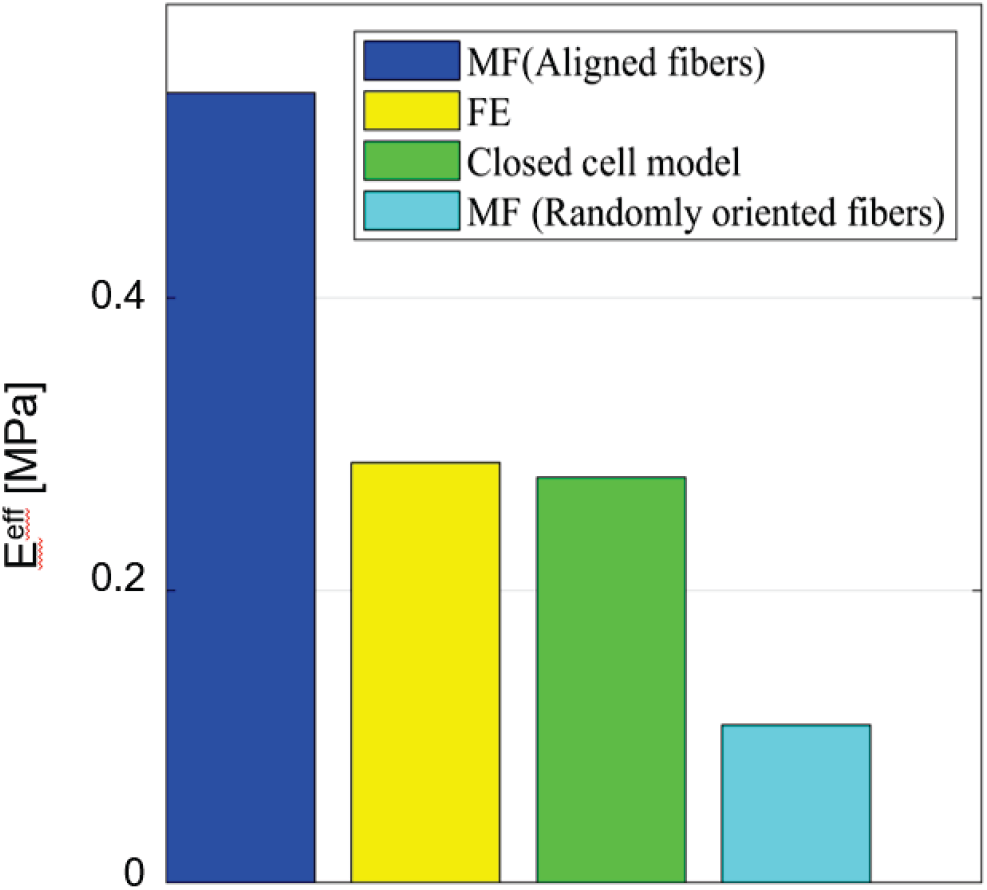
Values for the elastic modulus of the scaffold-hydrogel RVE obtained with different methods of homogenization.

### 3.2 Treatment effect of epicardial patching

The results of the LV in the healthy state, with untreated MI, and with MI treated with an epicardial patch show a slight improvement of stroke volume with epicardial patch compared to the untreated case, albeit lower than the healthy condition (Figure 8a). The patch provides mechanical restraint (i.e. limiting the infarct-related dilation at the end of passive filling) and elastic support (i.e. the stretched patch, stored with elastic energy at the end of passive filling, assisting the contractile function of the heart). Collectively, these effects resulted in an improved ejection fraction of 43% for patch treatment with elastic modulus of 0.287 MPa compared to 40% for the untreated infarct.

**Figure 8.**
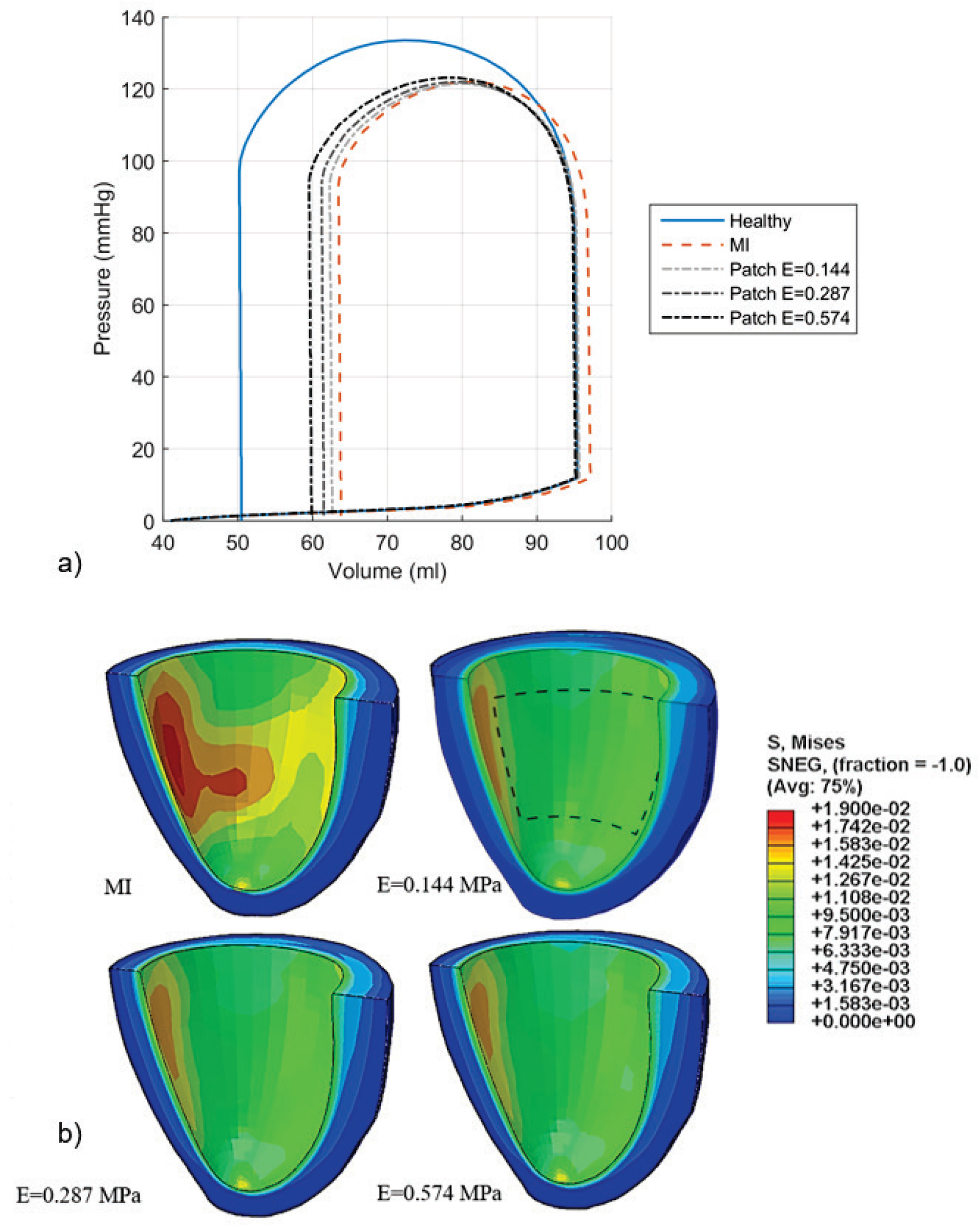
The application of an idealized patch to treat myocardial infarction improves function and regional stress within the heart. (A): Pressure volume loops comparing the cardiac output of the LV in a healthy state, with MI and with MI and patch treatment. (B): The stress distribution shown over half the LV revealing the endocardium surface of the “patched” region of the heart. The patch position is outlined in dashed lines.

The distribution and magnitude of the myocardial stress were also affected by the patch. High stresses experienced in the infarct region of the non-treated infarct case were not observed in the infarcted LV with patch treatment (Figure 8b), and the maximum value for myocardial stress decreased from 1.9e^−2^ MPa in the untreated infarct to 9.5e^−3^ MPa for the infarct treated with a patch with elastic modulus of 0.287 MPa.

## 4. Discussion

The fibrous scaffolds have been receiving attention for the use as cardiovascular tissue engineering patches [42]. Limitations of patches include, however, poor control of cell differentiation and unmatched mechanical properties [43]. Improving the understanding and control of the complex cell and tissue interactions in the scaffold and with the heart will enable the advancement towards clinical application.

In this study, we investigated the micro- and macro-mechanics of an electrospun polymeric scaffold filled with a hydrogel and the effect of such a scaffold as epicardial patch on myocardial mechanics and ventricular function in the presence of an acute myocardial infarct.

Our model of the scaffold was constructed from μCT images of a fibrous electrospun material. The hydrogel was assumed to entirely fill the void space between the fibres. Material properties for bulk polyurethane elastomer and a gentrified hydrogel were used [25], [26]. The model is based on simplified concepts of macro- and microscopic features of scaffold, hydrogel and cardiac loading. In the current simulation, the ventricular loading was simplified to maximal distention as specified in literature [44].

A global elastic modulus of the hydrogel-filled scaffold of 0.287 MPa was obtained from FE homogenisation. This value is set between upper (0.540 MPa) and lower limit (0.108 MPa) of the elastic modulus predicted by MF homogenization for completely aligned and randomly oriented fibres, respectively. Considering that the electrospun scaffold exhibited partially aligned fibres (Figure 1a), and the close agreement with the result of the closed cell model, verifies our image-based computational homogenisation method and results.

The simulation of the LV with MI demonstrated that the application of the epicardial patch, with material parameters derived through homogenization, resulted in somewhat improved stroke volume and reduced the peak stress values due to MI in the infarcted region in an acute setting. This in turn will have long-term benefit to the recovery and remodelling of the heart following MI [45], [46].

The FE homogenisation method provides a basis to guide epicardial patching in mechanical computational cardiac models of MI to optimise therapeutic effects. The approach can provide a computational framework for systematic screening of combinations of hydrogels and electro-spun fibres. The same framework could be used in a different way by running multi-objective design optimization studies, that is, a numerical optimiser would seek the structural and mechanical properties of the individual phases of the composite patch that lead to its optimal mechanical properties for patient-specific treatment.

This simplified model illustrates the utility of homogenization techniques that allow complex multiscale features to be represented in a simplified, yet meaningful manner. In our case, we provide mechanical insight into a potential therapy for MI. This is primarily meant as an illustration of the principle; further validation and sensitivity analysis is needed before this model could provide clinical insight.

## 5. Conclusion

Our current work intended to be a first step in *in silico* investigations. We sought to relate mechanical properties of a tissue-engineering scaffold across tissue and organ length scales. The analytical method and computational models developed for the micromechanical honogenisation of a composite structure offer potential for application to other areas in tissue engineering and regenerative medicine, offering both tailored mechanical support through the structural scaffold and biological effects through controlled and targeted drug release from the hydrogel. Future work will focus on extension towards cellular and sub-cellular length scales to link cellular mechanics into to the computational framework.

## Acknowledgements

The research reported in this publication was supported by the National Research Foundation of South Africa (UID 92531 and 93542), and the South African Medical Research Council under a Self-Initiated Research Grant (SIR 328148). Views and opinions expressed are not those of the NRF or MRC but of the authors. Nicola M. Pugno was supported by the European Commission with the Graphene Flagship Core 2 No 785219 (WP14 “Composites”) and FET Proactive “Neurofibres” No. 732344 as well as by the Italian Ministry of Education, University and Research with the “Departments of Excellence” grant L. 232/2016 and ARS01-01384-PROSCAN.

## Data

Abaqus models of the scaffold and the left ventricle used in this study are available on ZivaHub at http://doi.org/10.25375/uct.12894872.

